# Assessing neurocognitive maturation in early adolescence based on baby and adult functional brain landscapes

**DOI:** 10.1101/2024.09.26.615215

**Authors:** Omid Kardan, Natasha Jones, Muriah D. Wheelock, Cleanthis Michael, Mike Angstadt, M. Fiona Molloy, Lora M. Cope, Meghan M. Martz, Katherine L. McCurry, Jillian E. Hardee, Monica D. Rosenberg, Alexander S. Weigard, Luke W. Hyde, Chandra Sripada, Mary M. Heitzeg

## Abstract

Adolescence is a period of growth in cognitive performance and functioning. Recently, data-driven measures of brain-age gap, which can index cognitive decline in older populations, have been utilized in adolescent data with mixed findings. Instead of using a data-driven approach, here we assess the maturation status of the brain functional landscape in early adolescence by directly comparing an individual’s resting-state functional connectivity (rsFC) to the canonical early-life and adulthood communities. Specifically, we hypothesized that the degree to which a youth’s connectome is better captured by adult networks compared to infant/toddler networks is predictive of their cognitive development. To test this hypothesis across individuals and longitudinally, we utilized the Adolescent Brain Cognitive Development (ABCD) Study at baseline (9-10 years; n = 6,489) and 2-year-follow-up (Y2: 11-12 years; n = 5,089). Adjusted for demographic factors, our anchored rsFC score (AFC) was associated with better task performance both across and within participants. AFC was related to age and aging across youth, and change in AFC statistically mediated the age-related change in task performance. In conclusion, we showed that a model-fitting-free index of the brain at rest that is anchored to both adult and baby connectivity landscapes predicts cognitive performance and development in youth.

## 1. Introduction

Adolescence is a period of considerable brain development [Paus, 2005; Fuhrmann et al., 2015; Bethlehem et al., 2022], with noticeable changes in the landscape of the brain’s functional interactions (i.e., the functional ‘connectome’) [Zuo et al., 2017]. Understanding how variability observed in an individual’s connectome relates to their cognitive performance (e.g., sustained attention ability, working memory capacity), has been the subject of many recent studies [Rosenberg et al., 2016; Gao et al., 2019; Avery et al., 2020; Sripada et al., 2020]. However, relations between a brain phenotype and an outcome are likely to change across development [Cicchetti & Toth, 2009; Hyde et al., 2024], making it difficult to justify using age-agnostic static patterns in the connectome as predictors of cognitive performance and its development over adolescence [Kardan, Stier, et al., 2022a]. As such, methods to summarize the high-dimensional human connectome into age-dependent brain phenotypes that predict cognition are needed. This study aims to introduce and test a theoretically principled, interpretable, and simple to compute brain functional connectivity summary measure that captures cognitive performance and development during adolescence without the need for complex, data-driven prediction models.

In the past decade, data-driven methods utilizing machine learning have become widely popular in brain-behavior association studies [e.g., Maglanoc et al., 2020; Nielsen et al., 2020; Chen et al., 2022]. These methods provide multivariate solutions that are more reliable than univariate ones when modeling the connectome’s relation with behavioral measures [Yoo et al., 2019]. As an example, the “brain age gap” [Franke et al., 2010], which uses machine learning to identify a “neurobiological age” which is subtracted from chronological age (i.e., brain age gap), has been interpreted as a biomarker of deviations from normative aging and tied to mental health or cognitive decline [e.g. Biondo et al., 2022]. Recent work has started to extend this approach from aging populations to adolescents in cross-sectional and longitudinal studies [Holm et al., 2023; Dehestani et al., 2023; Whitmore et al., 2023; Rudolph et al., 2017]. Importantly, however, the utility of brain age gap in delineating cognitive development during adolescence is limited, with studies finding inconsistent or no relationships between brain age gap and cognitive performance [Erus et al., 2015; Lewis et al., 2018; Ball et al., 2021; Whitmore et al., 2023].

Another recently proposed connectomic measure for indexing cognitive maturation during adolescence is the degree of principality for primary-to-transmodal regions when decomposing the connectivity matrix [Margulis et al., 2016]. Briefly, gradient decomposition of the young children’s connectome reveals a somatosensory/motor-to-visual (i.e., unimodal) principal gradient. In adults, however, the unimodal gradient is secondary to a more dominant sensorimotor-to-default mode network regions (i.e., transmodal) gradient instead. Thus, the degree to which this shift has progressed has been proposed to track brain maturation and thus cognitive maturation in adolescence [Dong et al., 2021]. Associations between the primary-to-transmodal gradient and working memory performance have been reported *across* youth [Xia et al., 2022]. Another related index utilized recently to estimate the excitation-inhibition ratio in the cortex within age ranges covering adolescence have found similar results [see Zhang et al., 2024]. However, to our knowledge, no relations have been reported between the amount of change in either of these measures and the amount of cognitive development during adolescence.

Therefore, in the current study, we used a different approach to relate the inter- and intra-individual differences in cognitive performance of youth to their functional connectome. The central idea for our proposed index is that the cortex’s functional landscape (i.e., boundaries of different systems or networks) during childhood and adolescence periods is in transition from the early life brain systems to the mature brain networks [e.g., see Tooley et al., 2022]. Infancy and adulthood anchor the two points of this normative shift. If this general transition accommodates the cognitive maturation within the person, then the relative position of their connectome on this continuum should reflect their cognitive development. Specifically, more distance from the canonical baby functional landscape and less distance from the canonical adult functional landscape in an adolescent individual should predict better (i.e., closer to adult-like) performance compared to their peers. This hypothetical relationship is shown in **Figure 1**.

**Figure 1.**
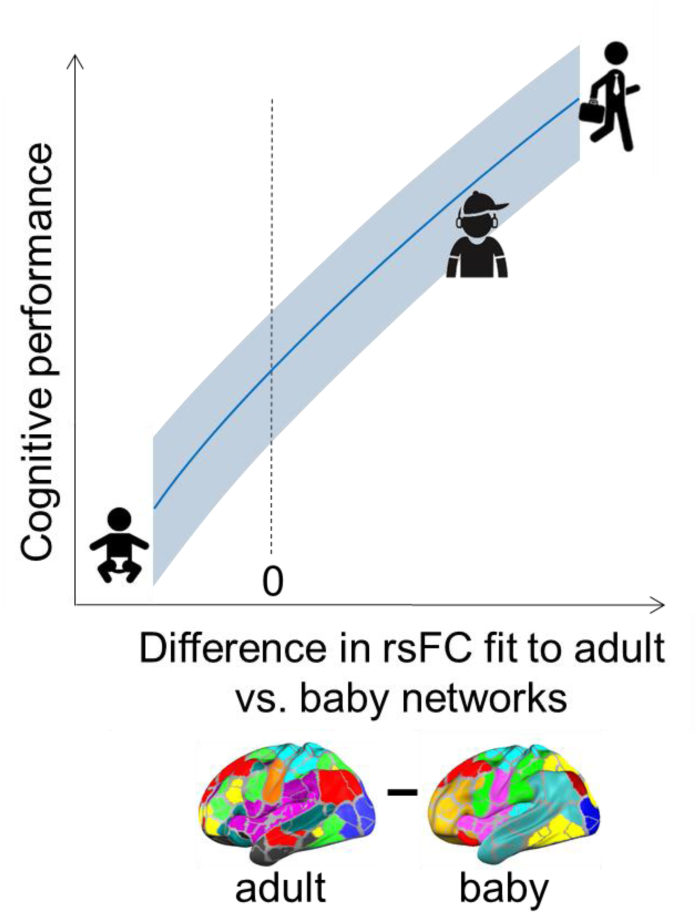
Hypothesized relationship between cognitive performance and anchored rsFC maturation score. The curve shows the relationship between cognitive maturation and the fit of resting-state functional connectome (rsFC) to adult vs. baby functional networks from infancy into adulthood (blue shade around the curve represents variability across individuals). The investigated regime in the current study (pre- to early adolescence) is shown with the icon with a baseball cap.

Our measure is not based on data-driven models that are trained to predict either age or cognitive performance from brain data and does not require two different sets of adolescent datasets for discovery and cross-validation. Our hypothesis is that the degree to which a youth’s connectome is better captured by adult networks compared to infant/toddler networks is predictive of their cognitive functioning and its development. To test this hypothesis, we utilized functional Magnetic Resonance Imaging (fMRI) data during rest of youth participating in the Adolescent Brain Cognitive Development (ABCD) Study. A participant’s matrix of connectivities between pairs of cortical regions was constructed both according to canonical baby cortical networks arrangement [Kardan et al., 2022] and canonical adult cortical networks arrangement [Gordon et al., 2016]. The fit to each of the baby or adult network assignments was measured with a simple index of mean of connectivities within networks minus mean of connectivities outside of networks (i.e., between networks) (see e.g., [Chan et al., 2014; Chan et al., 2018] for other works subtracting between-from within-network connectivities). The difference between these two measures of fit, henceforth referred to as the <u>a</u>nchored rs<u>FC</u> maturation score (AFC), provides a metric of neoteny vs. maturity of one’s connectome. We then tested whether AFC predicts performance in the N-back task across and within participants, and mediates the association between age and task performance.

In summary, in this study we propose a brain phenotype that is developmentally interpretable and is sensitive to both inter-individual and intra-individual differences in cognitive performance, thus addressing some of the current gaps in tracking neurocognitive maturation in this important period.

## 2. Methods

### 2.1. Participants

The Adolescent Brain Cognitive Development (ABCD) Study® is an ongoing longitudinal study of 11,867 children across 21 sites in the US [Casey et al., 2018]. Participants were enrolled in the study between 9–10 years of age, and the study involves MRI acquisitions every two years as well as behavioral assessments. The starting sample was 52.2% male and participants reported the following identities: 52.0% non-Hispanic White, 15.0% Black, 20.3% Hispanic, 2.1% Asian, and 10.5% Other.

We utilized the ABCD Study data Release 5.0 including baseline (Y0; ages 9-10) and Year 2 follow-up (Y2; ages 11-12) resting state fMRI and N-back task performance measures. After exclusions based on MRI data quality (see section 2.2), 6,604 and 5,168 participants remained for Y0 and Y2, respectively. Additionally, after exclusions based on missing N-back task or covariates, 6,489 and 5,089 participants remained for the main analyses at Y0 and Y2, respectively. In the longitudinal analysis, N = 3200 participants with complete data at both Y0 and Y2 were included. The demographic breakdown of these subsamples are shown in Table 1.

**Table 1.** Demographic breakdown of the included participants from the ABCD Study in the baseline (Y0), Year 2 follow-up (Y2), and the longitudinal (Y0 &Y2) analyses.

|  | Analysis Timepoint |  |  |
| --- | --- | --- | --- |
|  | Y0 | Y2 | Y0 & Y2 |
| Male | 3146 (48.5%) | 2578 (50.7%) | 1581 (49.4%) |
| Female | 3343 (51.5%) | 2511 (49.3%) | 1619 (50.6%) |
| White | 3750 (57.8%) | 2887 (56.7%) | 1961 (61.3%) |
| Black | 777 (12.0%) | 612 (12.0%) | 317 (9.9%) |
| Hispanic | 1180 (18.2%) | 980 (19.3%) | 576 (18.0%) |
| Asian | 131 (2.0%) | 88 (1.7%) | 47 (1.5%) |
| Other | 651 (10.0%) | 522 (10.3%) | 299 (9.3%) |

Additional exclusions based on task performance or family members were applied in sensitivity analyses and can be seen in Supplementary sections 1 and 2.

### 2.2. fMRI processing pipeline and exclusions

Resting-state fMRI data were acquired across four separate runs (∼5 min per run) for both Y0 and Y2 neuroimaging sessions (details are described in Hagler et al., 2019). The entire data pipeline was run through automated scripts on the University of Michigan’s high-performance cluster and is described in detail in [Sripada et al., 2021]. Images were acquired at temporal resolution of TR = 800 ms and spatial resolution of 2.4 mm isotropic. The processing pipeline included FreeSurfer normalization, ICA-AROMA, denoising, CompCor correction, and censoring of high motion frames with a 0.5 mm framewise displacement threshold. Visual quality control (QC) was conducted to assess registration and normalization steps. For the cross-sectional analyses, participants with at least two resting-state runs with each run having more than 250 degrees of freedom left in the BOLD timeseries after confound regression and censoring were included at either Y0 or Y2 (i.e., > 6.6 minutes total). For longitudinal analyses, those who met the above criteria at both baseline and Y2 were included.

### 2.3. Constructing functional connectivity matrices

For the rsFC analyses, resting-state fMRI data were spatially down-sampled to 333 parcels in the Gordon parcellation for cortical regions [Gordon et al., 2016]. Pearson correlations between activities in all pairs of these parcels were arranged in a 333*333 functional connectivity matrix for each participant at each imaging session (i.e., Y0 and Y2). Every connectome was arranged according to both a baby networks assignment and to an adult networks assignment, all defined in the same parcel space. Briefly, the adult networks atlas contained 12 networks that had been defined using Infomap community detection algorithm [Rosvall et al., 2008] in resting-state fMRI data from 108 young adults aged 18-33 [Gordon et al., 2016]. The baby networks atlas contained 10 networks defined with Infomap in resting-state fMRI data from 96 infants/toddlers from the Baby Connectome Project study [Howell et al., 2019] aged 9-24 months old [Kardan et al., 2022]. More details about the network assignments can be found in [Gordon et al., 2016] and [Kardan et al., 2022]. The anatomical maps of these network assignments are shown in the plots above diagonals of **Figure 2** (left panel for baby networks and right panel for adult networks), and can be seen in larger size in **Supplementary section 3.**

**Figure 2.**
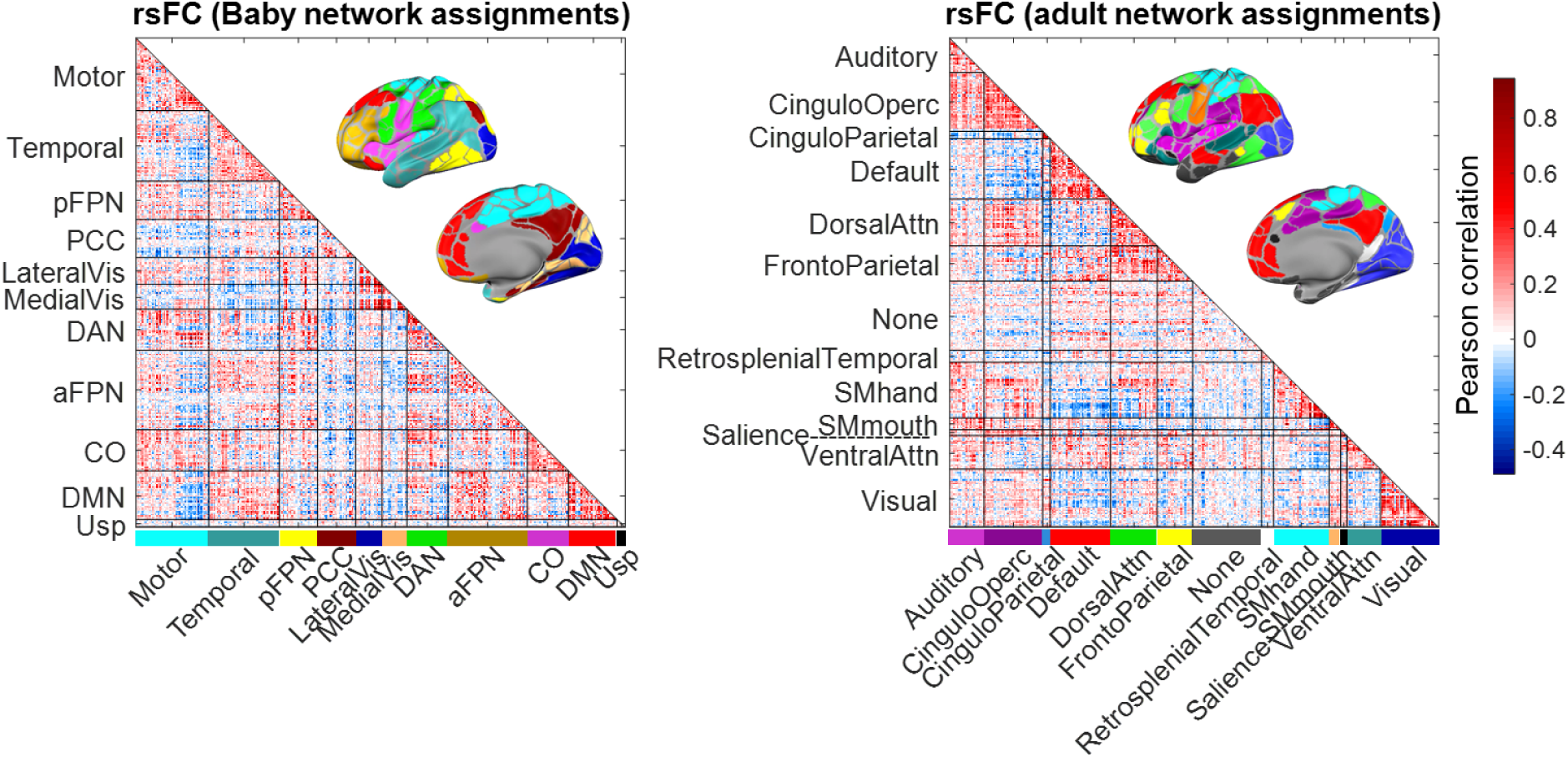
Example individual’s connectome arranged according to baby and adult network assignments. Above diagonals: The anatomical maps of the adult [Gordon et al., 2016] and baby [Kardan et al., 2022] networks assignments. **Below diagonals:** Demonstrating the calculation of the anchored rsFC maturation (AFC) score in a random participant’s connectome at age 11. In this participant, the difference in the Fisher Z-transformed mean connectivity of within-network and between-network connections is higher for the adult compared to baby network definitions ([*Conn*_*within*_ − *Conn*_*between*_]^*Adult*^ = 1.19 and [*Conn*_*within*_ − *Conn*_*between*_]^*Baby*^ = .79). The AFC score is the difference between the adult and baby values AFC = 1.19 - .79 = .40 in this example participant. Note: aFPN = anterior fronto-parietal network; pFPN = posterior fronto-parietal network; PCC = posterior cingulate cortex; Vis = visual cortex; CO = cingulo-opercular; DMN = default mode network; Usp and None both refer to unspecified remaining parcels. See Supplementary section 3 for full/other names of networks from [Gordon et al., 2016] and [Kardan et al., 2022].

### 2.4. Variables

#### Anchored rsFC maturation score

An individual connectome’s difference in within-network and between-network connectivity (henceforth shown as [Conn_within –_ Conn_between_]) was calculated as the mean of the Fisher Z transformed signed correlation values between pairs of parcels that were inside networks (Conn_within_) and those outside networks (Conn_between_). The [Conn_within –_ Conn_between_] is a measure of the functional adjacency of cortical parcels assigned to being part of a network compared to their connections to parcels outside [Kardan et al., 2022]. When averaged across all the networks of a certain atlas of functional networks, [Conn_within –_ Conn_between_] provides a simple measure of fit of a connectome to the assigned networks in an atlas. The [Conn_within –_ Conn_between_] was then calculated for each of the two networks assignments (i.e., baby and adult) and anchored rsFC maturation score was defined as:

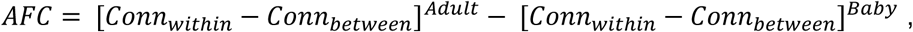

yielding one value for the connectome at Y0 and one value for the connectome at Y2 for each participant.

An example random participant’s cortical connectivity matrix arranged according to the two network assignments is shown in **Figure 2** below diagonals (i.e., lower traingles). The arrangement according to the baby network (left) or adult network (right) definitions shows that the adult networks better capture this rsFC matrix than the baby networks as more strongly positive connections are more along the diagonal (i.e., assigned as within-network) compared to off diagonal (i.e., assigned as between-network). The anchored rsFC score in this example connectome is +0.40. If the baby networks assignment better capture a connectome, then the value would be negative.

We also generated two other sets of network assignments to serve as null models. In the first null model, a set of random connections were assigned as within-network connections and [*Conn*_*within*_ − *Conn*_*between*_]^*Random*^ was calculated for each participant for these random networks. The total number of connections within the random networks matched the adult network sizes and the process was repeated 100 times for each participant. The second set of null networks were defined using only physical distance proximity of parcel centroids.

Specifically, k-means algorithm was used on the volumetric coordinates of the parcel centroids to partition the 333 cortical parcels into 12 clusters (or proximity-based null networks), where the parcels had minimum Euclidean distance from the center of their assigned cluster. The number of clusters was set to 10 or 12 to mimic the baby or adult networks numbers, but the produced null distributions were very similar, so the results reported here are based on the version with 12 clusters (see **Supplementary section 3** for visualization of these clusters). This null model would provide a more stringent and less naïve set of ‘networks’ than the random networks but still does not use the functional interactions of the parcels. Therefore, network assignments based on functional connectivity (baby or adult) should capture the connectome better than this proximity-based set of networks. [*Conn*_*within*_ − *Conn*_*between*_]^*Proximity*^ was then calculated for each connectome of each participant to serve as another null distribution.

#### Behavioral measures and covariates

Cognitive performance for each youth at Y0 and Y2 was measured as the accuracy in performing the N-back task [Casey et al., 2018]. This task taxes both sustained attention (0-back task condition) and working memory (2-back task condition) abilities, both of which show marked increase from pre-adolescence to adulthood (e.g., see Kardan, Stier, et al., 2022). Briefly, in each trial, participants had to indicate whether a picture is a ‘match’ or ‘no match’ to the picture either shown in the beginning of the block (0-back) or shown 2 trials prior (2-back). This task was performed in the scanner during two runs of 8 blocks each (160 trials total), but the task fMRI data were not used in this study.

Demographic and family socioeconomic status (SES) variables were extracted from the abcd_p_demo, ph_p_pds, ph_y_pds, and ph_y_anthro tables of the ABCD Release 5.0 files including parents reports on the participant’s biological sex, interview age, race/ethnicity, parental education (highest education), household income, and youth’s height and pubertal status (average of youth and parental reports on the Pubertal Developmental Scale). Race/ethnicity was controlled for to account for differences in exposure to personal/systemic racism, disadvantage, and opportunity among people of color. Family ID was used in supplementary analyses where only one sibling from each family was randomly retained (see supplementary results section 1). See shared scripts at https://github.com/okardan/Anchored_rsFC_maturation for more details of pulling these variables. Other covariates included head motion in the scanner during the resting-state fMRI (mean frame displacement), number of fMRI runs, as well as data collection site.

#### Connectome modularity

In our post-hoc analyses, we also compared the AFC score with a conceptually similar measure from graph theory, modularity. The rsFC matrices were constructed as weighted undirected networks and their community structure and modularity was quantified using the Newman’s spectral community detection [Newman 2006] similar to the implementation in Brain Connectivity Toolbox [Rubinov & Sporns, 2010].

### 2.5. Statistical analyses

All regressions were performed in R using the linear mixed-effects regression function lmer from the lme4 package. The mediations were performed using custom R functions utilizing lmer with confidence intervals estimated by bootstrapping. All scripts are available at https://github.com/okardan/Anchored_rsFC_maturation.

## 3. Results

### 3.1. Both adult and baby network assignments capture youth connectomes better than chance

We first established that both baby and adult network definitions fit youths’ brain functional landscape better than chance in at Y0 (ages 9-10) and also at Y2 (ages 11-12). This is an important data check, because the rsFC anchored maturation score is calculated as the difference between the fit of the youth’s connectome to the adult compared to baby networks (see section 2.2.; fit here is defined as better aggregation of strong positive connections within communities compared to outside of them). If each of these two network definitions (i.e., adult and baby networks) are not capturing the functional communities to an acceptable level in the youth age range, applying them to the youths’ brain data would be inappropriate. We tested this by comparing the distributions of [Conn_within –_ Conn_between_] for adult and baby communities to null distributions of [Conn_within –_ Conn_between_] (see **Figure 3**), where [Conn_within –_ Conn_between_] denotes the difference between the sum of connections (Fisher Z with sign) within networks and the sum of the rest of connections (i.e., between networks). The null distributions were generated by either assigning random connections as being within or between networks (grey distributions), or by simply partitioning the cortex into 12 communities based purely on proximity of the parcel centroids (pink distributions).

**Figure 3.**
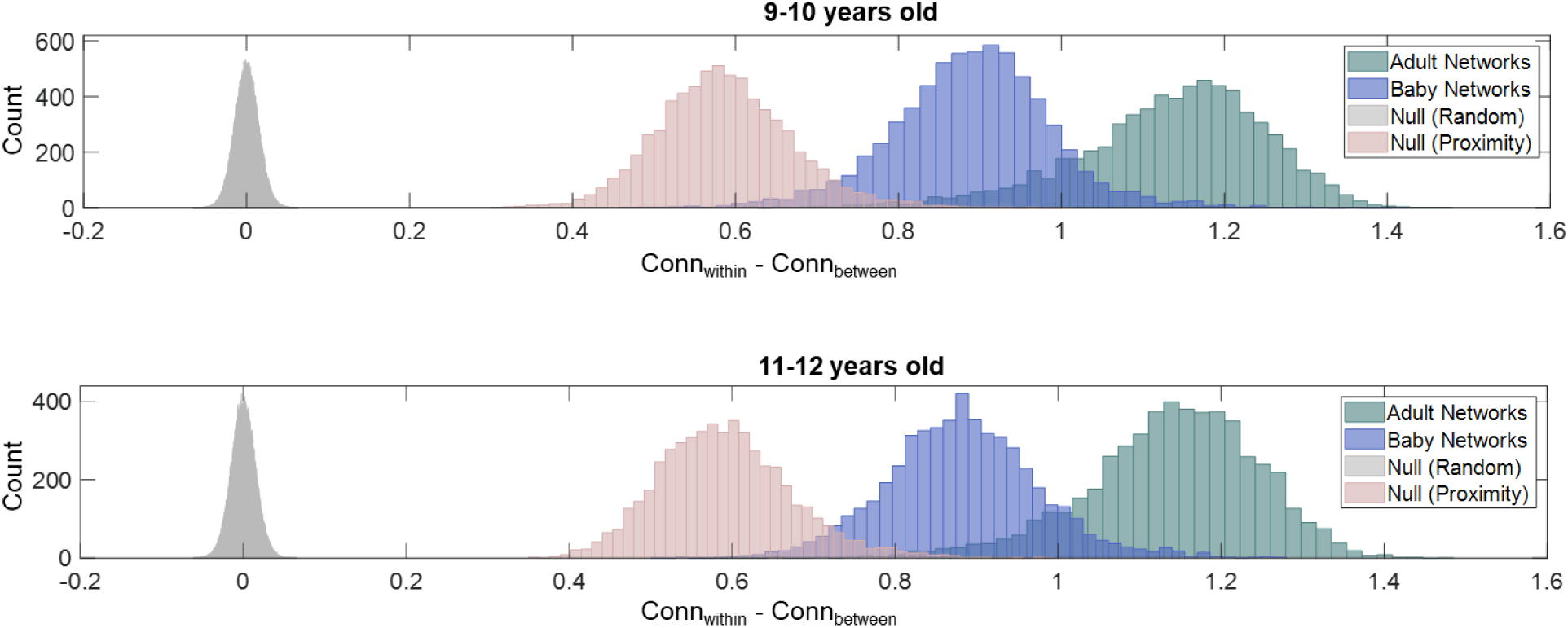
Assessing the fit of the assigned baby or adult communities to youths’ rsFC by comparing them to generated null communities (random or proximity-based). [Conn_within –_ Conn_between_] for a given participant is the difference between average Fisher Z-transformed connectivity of the edges within networks and the edges between networks in their connectome. The network assignments are based on adult communities (green; Gordon et al, 2014), baby communities (blue; Kardan et al., 2022), Euclidean distance proximity (pink), or completely random (grey). Top panel shows distributions for the ABCD baseline rsFC data and bottom panel shows them for the ABCD year two follow-up data.

As shown in **Figure 3** (top), at Y0, both baby and adult network assignments fit the youth connectomes better than the random and proximity-based network assignments. This is indicated by significantly stronger within-network compared to between-network connections than the proximity-based null networks (Cohen’s d = 3.39 for baby compared to proximity null networks; Cohen’s d = 5.57 for adult compared to proximity null networks; both ps < .001). This was also true at Y2 (**Figure 3**, bottom), with Cohen’s d = 3.28 for baby compared to proximity null networks and Cohen’s d = 5.79 for adult compared to proximity null networks (both ps < .001).

### 3.2. Anchored index of functional maturation predicts cognitive performance across and within youth

Next, we tested our hypothesis that the degree to which a youth’s connectome is better captured by adult networks compared to infant/toddler networks is predictive of their cognitive functioning and its development. The first set of analyses were adjusted for data collection site, the participant’s head motion, number of resting fMRI runs, sex, and race/ethnicity (but not parental income and education).

At both 9-10 and 11-12 years of age, we found that higher anchored rsFC maturation score was associated with higher N-back task performance across participants. **Figure 4A** shows the scatterplots for these relationships at Y0 (yellow; standardized adjusted β = .063, p < .001, N = 6,485) and Y2 (purple; standardized adjusted β = .099, p < .001, N = 5,089).

**Figure 4.**
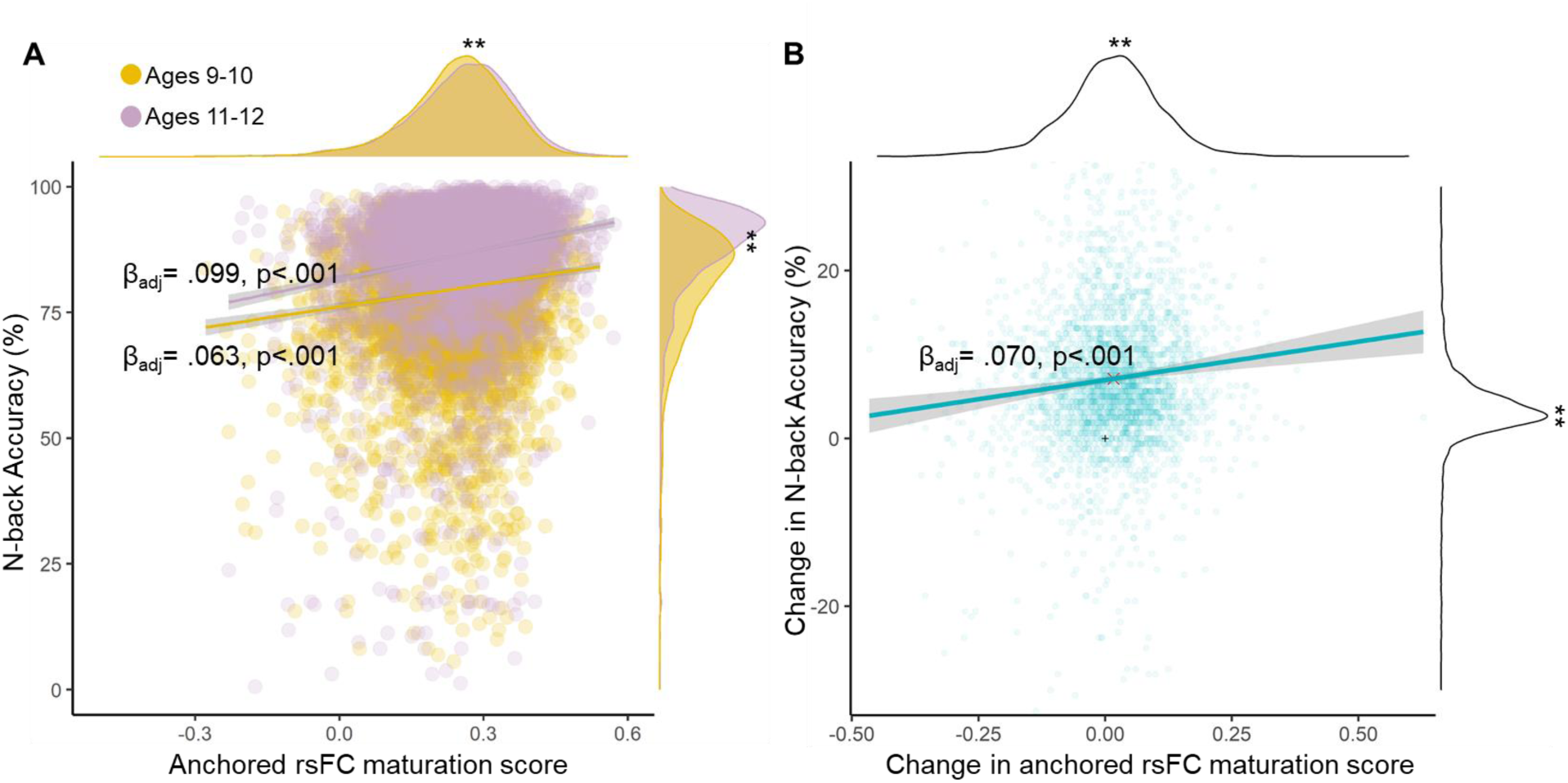
Anchored rsFC score predicts N-back accuracy across and within youth. **A.** maturation score was associated with higher N-back task performance across participants at baseline (yellow) and at two-year-follow-up (purple). Margins: ** indicates p < .001 for difference between the two distributions. **B.** The amount of change in the anchored rsFC score was associated with the amount of change in the N-back task performance from baseline to two-year-follow-up across participants. Margins: ** indicates p < .001 for difference between the distributions and 0.

Additionally, there was a significant increase in the anchored rsFC score from Y0 to Y2 both in aggregate (Welch two-sample t = 5.62, p < .001; **Figure 4A** distributions on the top), as well as within participants (paired t = 9.24, p < .001; **Figure 4B** distributions on the top). The amount of change from Y0 to Y2 in the anchored rsFC score (i.e., AFC_Y2_ – AFC_Y0_ or ΔAFC) was associated with the amount of change in the N-back task performance (i.e., Acc_Y2_ – Acc_Y0_ or ΔAccuracy) across the participants **(Figure 4B**, standardized adjusted β = .070, p < .001 without additionally adjusting for baseline N-back accuracy; β = .071, p < .001 with additionally adjusting for baseline N-back accuracy; β = .094, p < .001 with additionally adjusting for baseline N-back accuracy and baseline AFC; N = 3,200).

Furthermore, sensitivity analyses showed these results are robust to excluding participants whose accuracy in N-back task was lower than 60% (**Supplementary section 1**) or randomly retaining only one sibling from each family (**Supplementary section 2**).

### 3.3. AFC predicts cognitive performance above and beyond family resources

In the second set of analyses, we added parental education and income to the model. This step was separated (i.e., instead of having family SES as another covariate in the first analysis) to investigate potential relationships between the anchored rsFC score, and its growth, to parental education and income, both of which are tied to individual differences in adolescent brain development [Foulkes et al, 2018; Michael et al., 2024].

First, we found that, adjusted for data collection site, head motion, number of resting fMRI runs, sex, and race/ethnicity, higher parental education was associated with higher rsFC maturation score (standardized adjusted β = .044, p < .001 at Y0; β = .040, p < .001 at Y2; Note: income is included in these models). Higher income was also associated with higher anchored rsFC score, but only at Y2 (standardized adjusted β = .005, p = .299 N.S. at Y0; β = .042, p < .001 at Y2; Note: education is included in these models). Additionally, higher income was associated with a larger increase in rsFC maturation score from Y0 to Y2 (standardized adjusted β = .064, p < .001; education is included in the model).

Second, we found that the associations between anchored rsFC score and N-back task performance (i.e., βs from the analyses in **Figure 4A**) were not significantly changed when adding the income and education variables to the model (at Y0: median change in β = .007, bootstrap p = 0.107, N.S.; at Y2: median change in β = .002, bootstrap p = 0.347, N.S.). Additionally, the association between change in anchored rsFC score and change in N-back performance from Y0 to Y2 (i.e., β from **Figure 4B**) did not significantly change either (median change in β = .005, bootstrap p = 0.174, N.S.). In summary, anchored rsFC score was related to family resources, but predicted task performance independently. However, in all of the analyses that follow, income and education are included as additional covariates.

### 3.4. AFC as a statistical mediator of age’s association with cognitive task performance

Next, we investigated the relationships between the AFC and its growth (i.e., ΔAFC) to age and aging (i.e., Δage), as well as other youth maturation phenotypes (pubertal status and height). Although the two-year windows in age and aging in the datasets are relatively small, we proposed the anchored rsFC score with the intention of introducing a relatively sensitive neurocognitive maturation signature for the adolescence period. Therefore, these samples can provide a benchmark for assessing how much of the age-related cognitive maturation can be captured by this measure.

First, we found a significant association between anchored rsFC score and age, but not pubertal status or height, across youth at baseline (Y0; ages 9-10 years, standardized adjusted β = .038, p < .001, β is adjusted for site, head motion, number of resting fMRI runs, sex, race/ethnicity, as well as parental education and income). Interestingly, the association between anchored rsFC score and N-back task performance across participants remained significant after additionally adjusting it for age, pubertal status, and height (standardized adjusted β = .044, p < .001). However, this did lead to a significant decrease in the regression coefficient for anchored rsFC score predicting N-back accuracy (median change in standardized adjusted β = .007, bootstrap p = .003). Consequently, we ran an indirect effects model (i.e., performed a statistical mediation analysis) to test if anchored rsFC score statistically mediates the relationship between age and task performance across the participants at Y0. The results showed a significant indirect effect such that older youths’ higher performance in the N-back task was statistically mediated by their higher anchored rsFC score (See **Figure 5A**; p = .021 for the mediated effect β_a*b_ based on bootstrapping).

**Figure 5.**
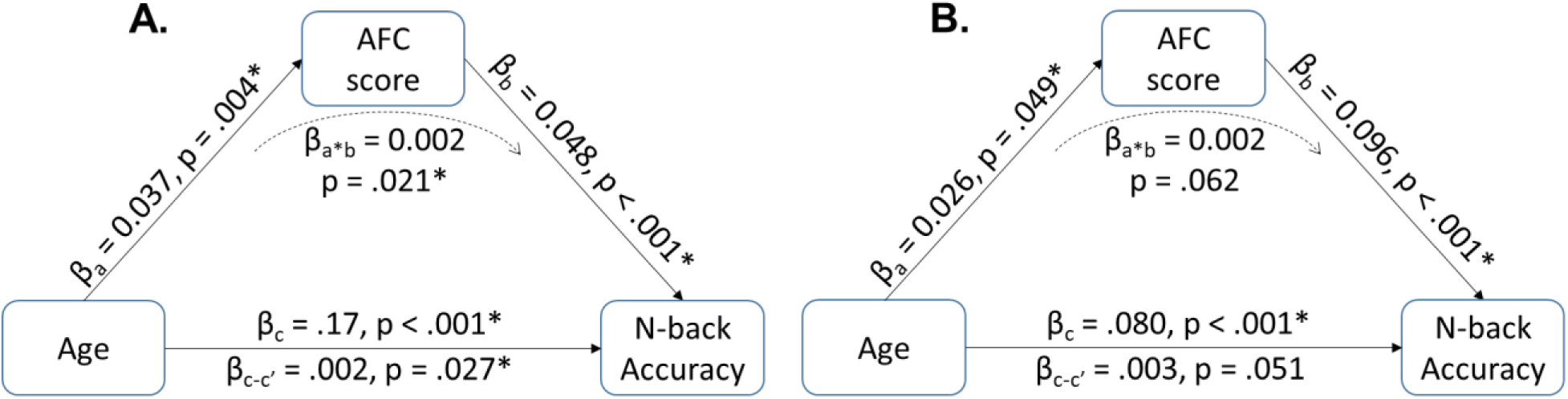
Anchored rsFC (AFC) score may statistically mediate the age’s association with cognitive performance in 9-10 and 11-12 years of age. **A:** Statistical mediation model at Y0. **B:** Statistical mediation model at Y2.

We then repeated the indirect effects model in the Y2 sample. We did not find anchored rsFC score to be a significant mediator of the association between age and task performance at Y2, although the indirect effect was trending in the expected direction (See **Figure 5B**; p = .062 for the indirect effect β_a*b_ based on bootstrapping). Interestingly, the association between task performance and anchored rsFC score was at least as strong as age (i.e., β_b_ is numerically larger than β_c_) in Y2 sample. Additionally, in both Y0 and Y2, older youth had higher anchored rsFC scores (β_a_ in **Figures 5A** and **5B**). All mediations include data collection site, head motion, number of resting fMRI runs, sex, race/ethnicity, parental education and income, pubertal status, and height as covariates.

We previously showed that AFC increases from Y0 to Y2 within participants and the amount of this change is associated with the amount of change in task performance (**Figure 4**). However, whether variation in the amount of Y0-to-Y2 age change between participants is also reflected in the variation in the amount of change in anchored rsFC score or not was not addressed. Therefore, we tested if Δage was associated with change in anchored rsFC score (ΔAFC) across the participants to supplement our cross-sectional indirect effects models reported above. It should be mentioned that, because the ABCD Study participants mostly returned for their Y2 fMRI scans relatively close to 24 months after Y0, the variance in age change (Δage) is low (mean Δage = 24.6 months; SD of Δage = 2.4 months).

First, we found that more increased age from Y0 to Y2 was associated with more increased AFC (standardized adjusted β = .037, p = .045), adjusted for baseline age, pubertal status, and height, change in pubertal status, change in height, as well as the other covariates from earlier models at both Y0 and Y2 (note: for site, family SES, sex, and race/ethnicity, only Y0 values were used due to little to no change). Above Δage, change in pubertal status was not significantly associated with change in anchored rsFC score (standardized adjusted β = .032, p = .182).

Second, an analysis with anchored rsFC score as mediator showed an indirect-only mediation [Zhao et al., 2010] between the amount of change in N-back task performance and Δage through the AFC (See **Figure 6**; p = 0.040 for the mediated effect β_a*b_ based on bootstrapping; effects are adjusted for all covariates mentioned above and baseline N-back accuracy).

**Figure 6.**
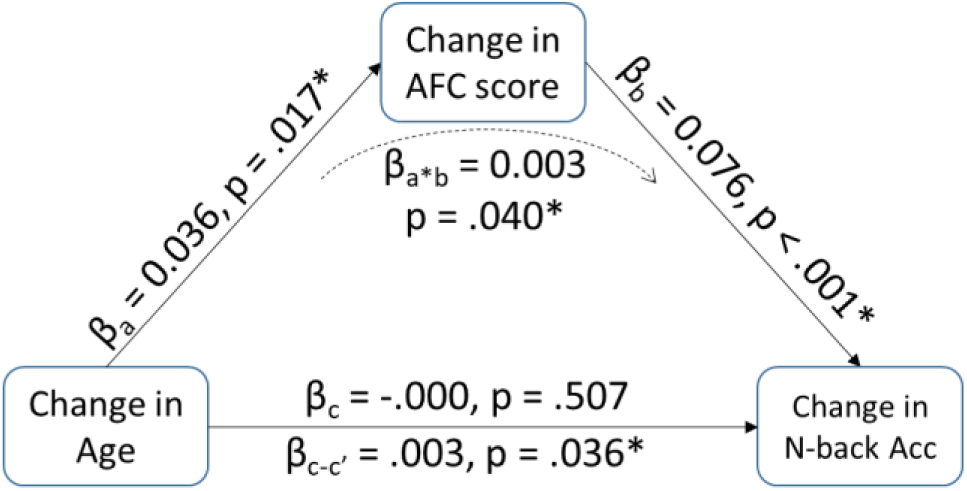
Anchored rsFC (AFC) score may mediate the effect of aging on cognitive development from 9-10 to 11-12 years of age.

In summary, we found evidence suggesting that anchored rsFC score is related to not only task performance across and within youth (section 3.2), but also their age (the β_a_s in Figure 5), aging (the β_a_ in Figure 6 and distributions in Figure 4A top), and age-related changes in task performance (the β_a*b_s in Figures 5 and 6).

### 3.5. Does anchoring to baby networks matter?

In the final section, we performed a series of post-hoc analyses to determine the degree to which anchored rsFC score truly provides a *novel* index of neurocognitive maturation in the brain in the adolescence period.

First, we compared the performance of the anchored rsFC score with another measure borrowed from graph theory that is commonly used in network neuroscience, brain modularity (e.g., [Cohen & D’Esposito, 2016]). The two measures are conceptually similar because they both relate to system segregation in the functional connectome. However, unlike the anchored rsFC score used here, modularity is quantified for the specific connectome of the participant based on communities that best capture that connectome, rather than based on previously constructed group normative communities.^1^ We found no significant associations between the connectome modularity and N-back accuracy across youth at Y0 (standardized adjusted β = - .020, p = .097; N = 6489) or at Y2 (standardized adjusted β = .002, p = .885; N = 5089). Additionally, change in modularity from Y0 to Y2 did not have a significant relationship with change in N-back accuracy from Y0 to Y2 (standardized adjusted β = -.002, p = .904; N = 3200). These indicated that anchoring to normative networks is paramount to construct our measure of neurocognitive maturation.

Second, measures approximating the ‘precocious’ development of functional brain based on <u>adult</u> brain architecture have been previously used in studies of cognitive development [Petrican et al., 2017]. Here, however, the idea of adolescence being sandwiched between baby and adult periods and therefore introducing distance from the <u>baby</u> networks into our measure is an important part of the novelty of our method. As such, we tested if anchoring to baby networks was indeed needed to delineate cognitive performance and development in youth, or if we would have achieved the same results had we simply used the adult networks. We found that the [Conn_within_ - Conn_between_] values for adult networks only (i.e., no subtraction of the baby networks value) did not have a significant relationship with N-back accuracy across participants at Y0 (standardized adjusted β = .014, p = .314; N = 6489). While, adult [Conn_within_ - Conn_between_] had a significant association with N-back accuracy at Y2 (standardized adjusted β = .032, p = .033; N = 5089), the strength of this association was significantly reduced compared to the AFC score (See **Figure 4**; median change in β = .077, bootstrap p < .001). On the other hand, baby [Conn_within_ - Conn_between_] did have a significant negative association with task performance at both Y0 and Y2 (standardized adjusted β = -.061, p = .001 at Y0; β = -.079, p = .001 at Y2), with the magnitude of βs being only numerically smaller than the AFC’s βs (at Y0 : median change in magnitude of β = .002, p = .450; at Y2 median change in magnitude of β = .031, p = .061).

Therefore, anchored rsFC score provides different information than brain network modularity, and its predictive power for cognitive performance in the age range of the current study lies more heavily in its early-life than adult component (i.e., the distance from baby networks rather than the proximity to adult networks).

## 4. Discussion

In this study we assessed neurocognitive maturation status in early adolescence by directly comparing a youth’s resting-state connectome to the canonical communities in early life and adulthood. Utilizing the large and heterogeneous ABCD Study sample at ages 9-10 and 11-12, we found that anchored functional connectome maturation score (AFC) was associated with cognitive performance across and cognitive development within youth. Additionally, change in AFC from 9-10 to 11-12 years was associated with higher family resources, but its relationship with cognitive development was above and beyond family resources. AFC was sensitive to age and aging, and mediated age’s relationship with task performance. Therefore, the current study proposes AFC as an unsupervised connectomic measure with theoretical and empirical links with cortical functional maturation that accommodates cognitive development.

AFC is easy to compute and apply to different studies/samples. Furthermore, utilizing an unsupervised approach confers other benefits. The majority of work using functional connectomes to characterize inter- and intra-individual differences in behavior has relied on supervised prediction methods (e.g., [Yeung et al., 2022]). Supervised approaches identify relevant connectome features, such as subsets of connections or graph theoretic properties, formalize the relationship between these features and a behavior of interest in a training set of data, and then apply the resulting model to forecast outcomes in novel individuals or datasets. In contrast, most theory-driven/unsupervised approaches relate a brain feature to behavior without the need for feature selection and model building in a training set. This has the benefit of maximizing sample size for detecting brain–behavior relationships and may confer benefits for predictive power and model generalizability. For example, recent work demonstrated that unsupervised models captured individual differences in attention and working memory even in datasets to which previously validated supervised connectome-based models failed to generalize [Corriveau et al., 2023]. Last but not least, many popular supervised models that work well across individuals provide only predictions of the participant’s standing relative to others (i.e., their rank order). Although this is well-justified in other fields, in developmental network neuroscience, magnitude of change over time may be meaningful and of interest even if rank stays unchanged. Our results show AFC increases longitudinally and the amount of change in AFC captures meaningful variability across youth in their trajectory of cognitive performance.

Although the associations we identified between AFC and cognitive performance were robust across ABCD Study waves and across levels of analysis (i.e., between- and within-person differences), the absolute value of the effect sizes for these associations is considered “small” (adjusted standardized βs ≤ 0.10) in the context of popular effect size benchmarks. In contrast, data-driven supervised models in cross-sectional data have been able to identify associations between distributed rsFC signatures and cognitive measures in ABCD that fall in the “moderate” to “large” range (*r* = 0.30 - 0.50) of such benchmarks [e.g., Sripada et al., 2020; Sripada et al., 2021; Marek, Tervo-Clemmens et al., 2022; Kardan, Stier, et al., 2022]. The smaller effect sizes yielded by AFC are to be expected given that the measure is based on a principled theoretical postulate about brain maturation rather than on a data-driven algorithm that is unconstrained by any theoretical assumptions and instead seeks to maximize variance explained. Data-driven approaches naturally maximize the effect sizes of associations but are difficult or impossible to interpret with regard to the mechanistic processes that drive the association [Huys et al., 2016; Yarkoni & Westfall, 2017]. Furthermore, it is important to note that most effects in brain-behavior association studies [Marek, Tervo-Clemmens et al., 2022] and even most effects between behavioral measures in large population-based samples like ABCD [Owens et al., 2021] also tend to fall in the “small” range by conventional standards. Indeed, it has been recently proposed [Funder & Ozer, 2019] that most plausible effects in psychological and biomedical research are likely to be in this range, and that such effects can have substantial practical consequences at the population level.

The baby [Kardan et al., 2022] and adult networks [Gordon et al., 2016] used in our study are both from data-driven resting state cortical parcellations defined using large samples of individuals within their respective age groups. Consistent with prior work in neonates and infants, our baby networks separate along higher order association cortex (i.e., DMN and frontoparietal separate on anterior posterior axis) [Wang et al., 2023; Eggebrecht et al., 2017; Sylvester et al., 2023; Eyre et al., 2021]. The [Gordon et al., 2016] atlas is widely used in studies with samples throughout development, but numerous other parcellations have also been proposed [Eickhoff et al., 2018]. While [Conn_within_ - Conn_between_] could be compared with any ROI set and any networks, the [Kardan et al., 2022] networks preserve the features (parcels) from the [Gordon et al., 2016]. This combined with the fact that the network assignments were computed in the same way (Infomap community detection) in both atlases makes the components of the AFC measure more comparable. However, important differences exist between infant and adult networks. Adult functional networks have been validated with task activation and myelination/cytoarchitectonic borders while infant networks generally have not been validated in these ways. Further, the infant networks are derived during sleep while the adult networks are derived from awake resting-state data which may also contribute to observed differences and poorer fit with ABCD Study awake rsFC. Despite these differences, AFC, as a unique metric of brain development and the neoteny of the functional connectome, seems to be an informative indicator of cognitive function by capturing important developmental information embedded within the network organization.

With regards to the pace of connectome development, on the one hand, adolescents whose network affiliations conform more closely to adult versus baby architectures may be further along in their neurocognitive development. On the other hand, other theories highlight that human brain development is protracted (even extending into adulthood), and that this slow developmental course may provide more time for the developing brain to “select” and “strengthen” the most relevant connections, while “pruning” away the least relevant connections, ultimately leading to more efficient and specialized brain systems (e.g., [Tooley et al., 2021]). This process may also be shaped by the environment afforded to each child. For example, different models propose that adversity and disadvantage may accelerate brain development [Callaghan & Tottenham, 2016; Tooley et al., 2021; Gee, 2021], although evidence for delayed brain development has also been reported [Johnson et al., 2016; Hyde et al., 2024]. In this study, there is evidence that adolescents whose brain development may be characterized as progressing faster exhibit better cognitive performance, and that this process may be scaffolded by greater access to socioeconomic resources. However, boundary conditions that shape whether and how pace of neurodevelopment may relate to cognition as a function of context remain poorly understood.

There are some limitations to the current study. First, the narrow age range of the included sample highlights the sensitivity of AFC, but also necessitates further assessments in mid-to-late adolescent populations. Second, during adolescence significant structural and functional brain changes also occur in the cerebellum and subcortical areas, but the networks used here only include the cortex. However, if we define cerebellum, basal ganglia, and thalamus clusters along their anatomical boundaries for the baby atlas, adding them to the cortical connectomes would have no impact on the AFC because the network definitions for subcortex and cerebellum would be identical to their adult counterparts for each participant. Future work with nuanced functional clusters specific to early life versus adult subcortical/cerebellum systems could expand the cortical AFC measure to whole-brain coverage. Last but not least, though the Y0 and Y2 samples matched population estimates for marginalized racial-ethnic groups (e.g., 12% African-American/Black), the Y0-to-Y2 longitudinal sample did not (e.g., 9.9% African-American/Black). Therefore, future work should assess AFC in longitudinal samples that are more representative of the US and worldwide populations.

In conclusion, in this study, we introduced and tested a theoretically principled and interpretable brain functional connectivity summary measure, anchored rsFC maturation score, that captures cognitive performance and development based on the neoteny of the youth’s functional connectome.

## Data and Code availability

Scripts to generate the results and figures in this study are available at https://github.com/okardan/Anchored_rsFC_maturation. Data tables used in the scripts can be downloaded from the https://nda.nih.gov/study.html?id=2147 after creating an account and being approved on an ABCD Study data use certificate.

## Acknowledgements

O.K. was supported by the National Institute on Drug Abuse K01 DA059598. M.F.M. was supported by National Institute on Alcohol Abuse and Alcoholism T32 AA007477. A.W. was supported by K23 DA051561.

## ABCD Acknowledgements

Data used in the preparation of this article were obtained from the Adolescent Brain Cognitive Development (ABCD) Study (abcdstudy.org), held in the NIMH Data Archive (NDA). This is a multisite, longitudinal study designed to recruit more than 10,000 children age 9 to 10 and follow them over 10 years into early adulthood. The ABCD Study is supported by the National Institutes of Health and additional federal partners under award numbers U01DA041022, U01DA041028, U01DA041048, U01DA041089, U01DA041106, U01DA041117, U01DA041120, U01DA041134, U01DA041148, U01DA041156, U01DA041174, U24DA041123, and U24DA041147. A full list of supporters is available at abcdstudy.org/nih-collaborators. A listing of participating sites and a complete listing of the study investigators can be found at abcdstudy.org/principal-investigators.html. ABCD consortium investigators designed and implemented the study and/or provided data but did not necessarily participate in analysis or writing of this report. This manuscript reflects the views of the authors and may not reflect the opinions or views of the NIH or ABCD consortium investigators. The ABCD data repository grows and changes over time. The ABCD data used in this report came from NIMH Data Archive Digital Object Identifier 10.15154/8873-zj65.

## Supplementary Material

### Exclusion of family members

We randomly retained only one sibling at Y0 and Y2 and repeated the analyses relating the inter- and intra-individual differences in cognitive performance to anchored rsFC score (N = 5,641 at Y0; N = 4,494 at Y2; N = 2,739 for Y0 & Y2).

**Figure S1.**
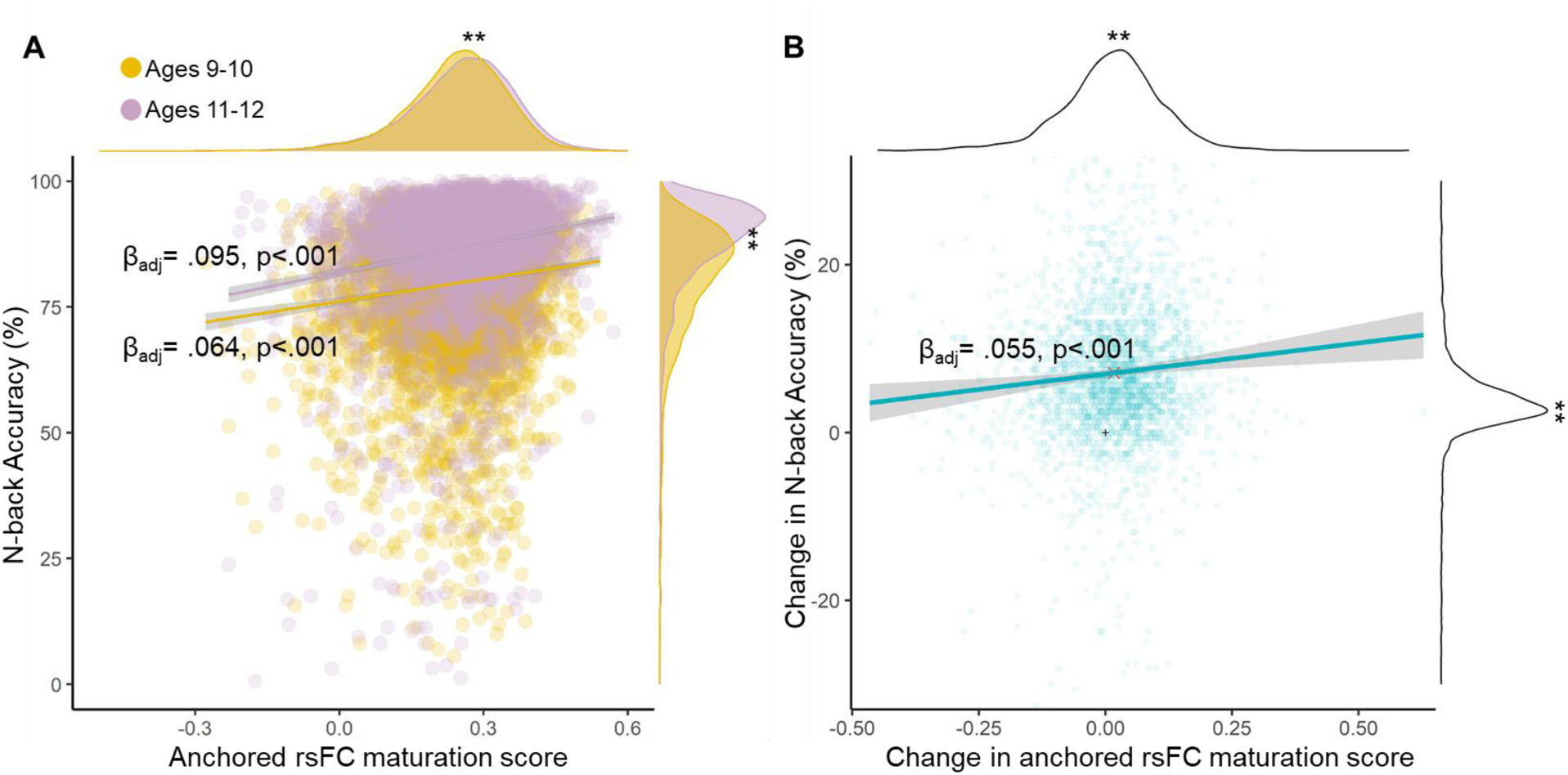
Anchored rsFC score predicts N-back accuracy across and within youth after including only one random sibling from each family. **A.** Maturation score was associated with higher N-back task performance across participants at baseline (yellow) and at two-year-follow-up (purple). Margins: ** indicates p < .001 for difference between the two distributions. **B.** The amount of change in the anchored rsFC score was associated with the amount of change in the N-back task performance from baseline to two-year-follow-up across participants. Margins: ** indicates p < .001 for difference between the distributions and 0.

### Exclusion of participants with low task accuracy

We excluded the participants with N-back accuracy lower than 60% at Y0 and Y2 and repeated the analyses relating the inter- and intra-individual differences in cognitive performance to anchored rsFC score (N = 6,060 at Y0; N = 4,924 at Y2; N = 2983 at Y0 & Y2).

**Figure S2.**
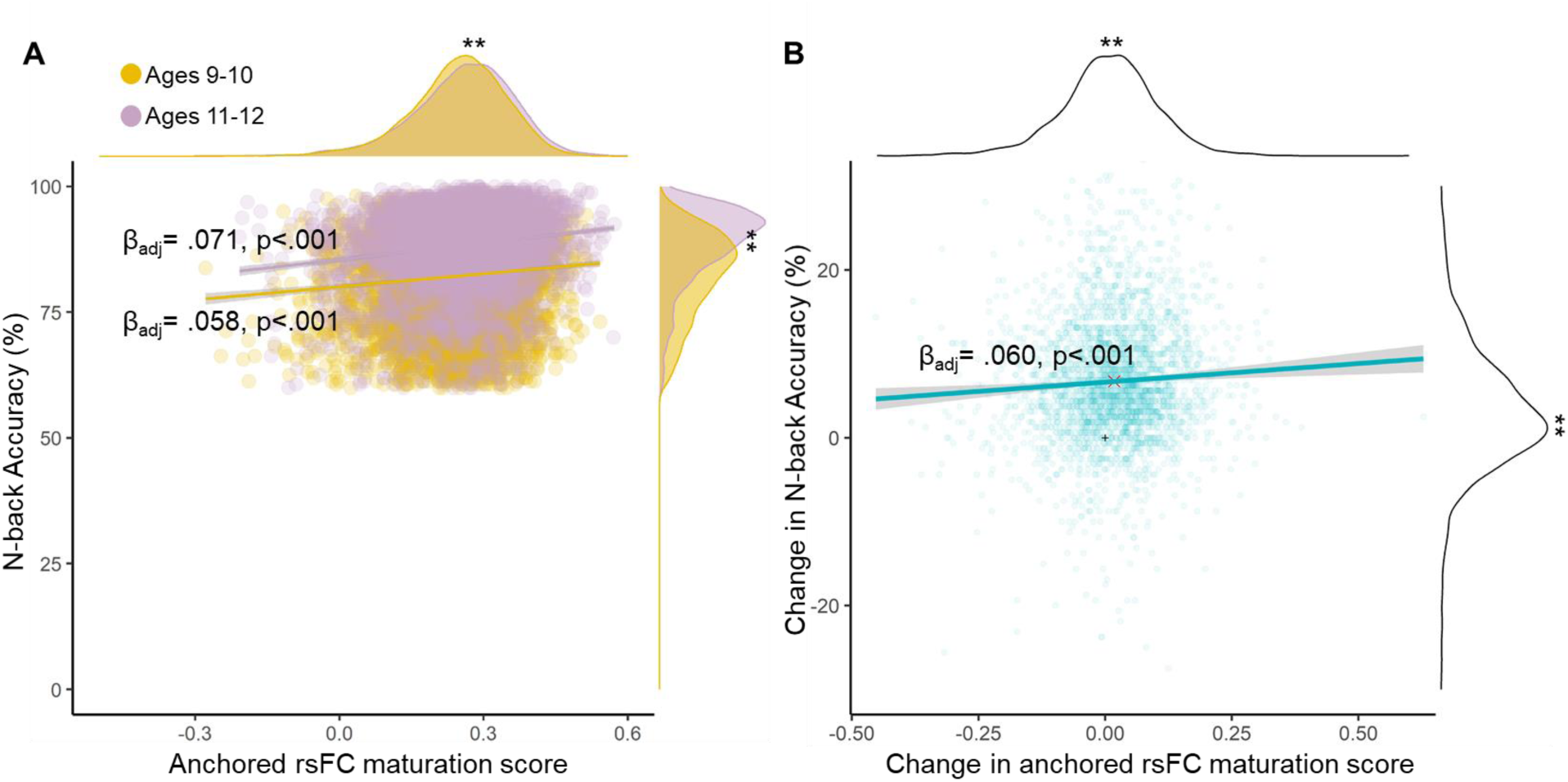
Anchored rsFC score predicts N-back accuracy across and within youth after excluding low-performing participants. **A.** Maturation score was associated with higher N-back task performance across participants at baseline (yellow) and at two-year-follow-up (purple). Margins: ** indicates p < .001 for difference between the two distributions. **B.** The amount of change in the anchored rsFC score was associated with the amount of change in the N-back task performance from baseline to two-year-follow-up across participants. Margins: ** indicates p < .001 for difference between the distributions and 0.

### Adult and baby networks assignments and the proximity-based null networks

**Figure S3.1.**
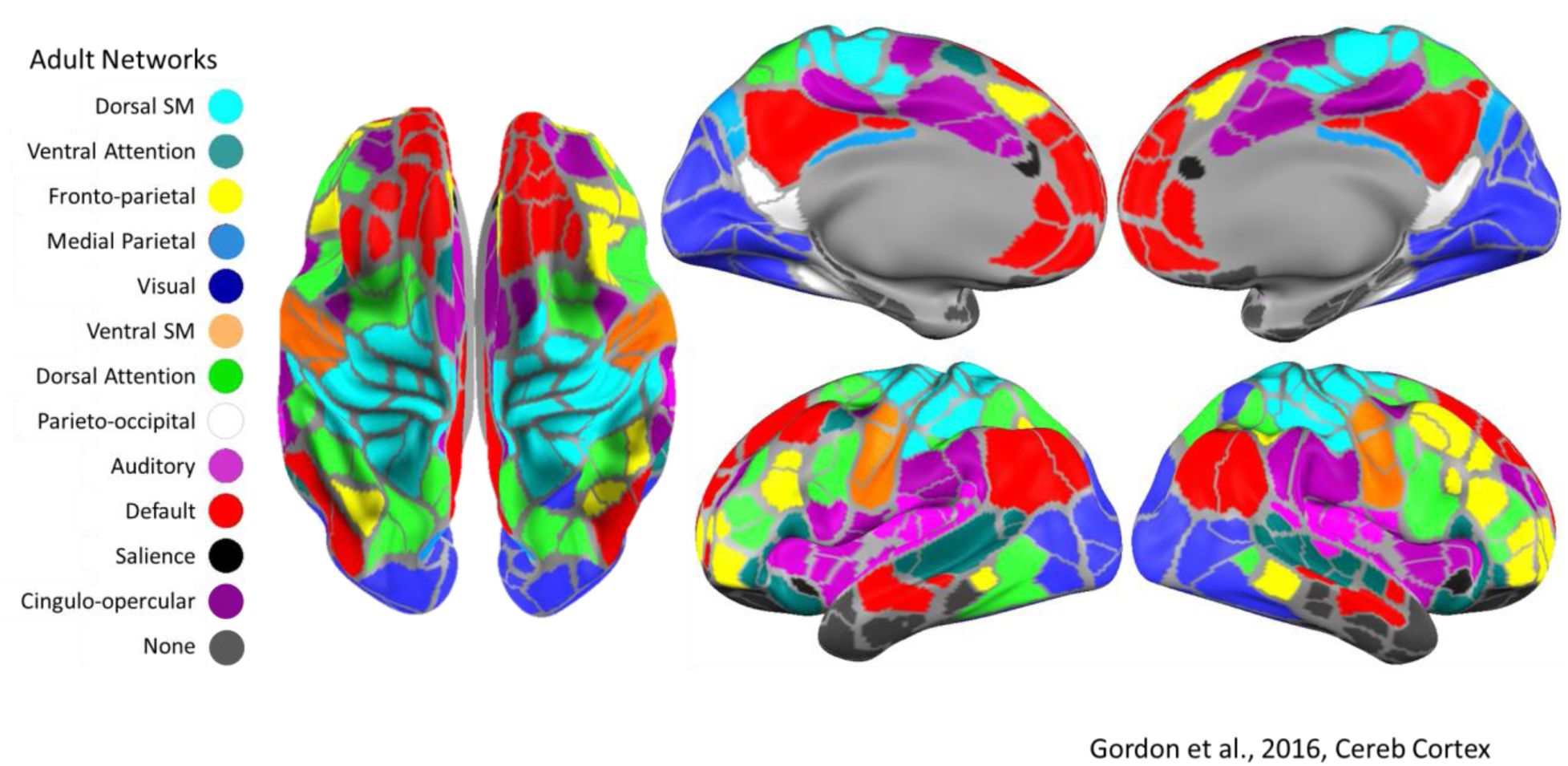
Adult network assignments. Note: Dorsal SM = Dorsal Somato-Motor (also SM-hand); Ventral SM = Ventral Somato-motor (also SM-mouth)

**Figure S3.2.**
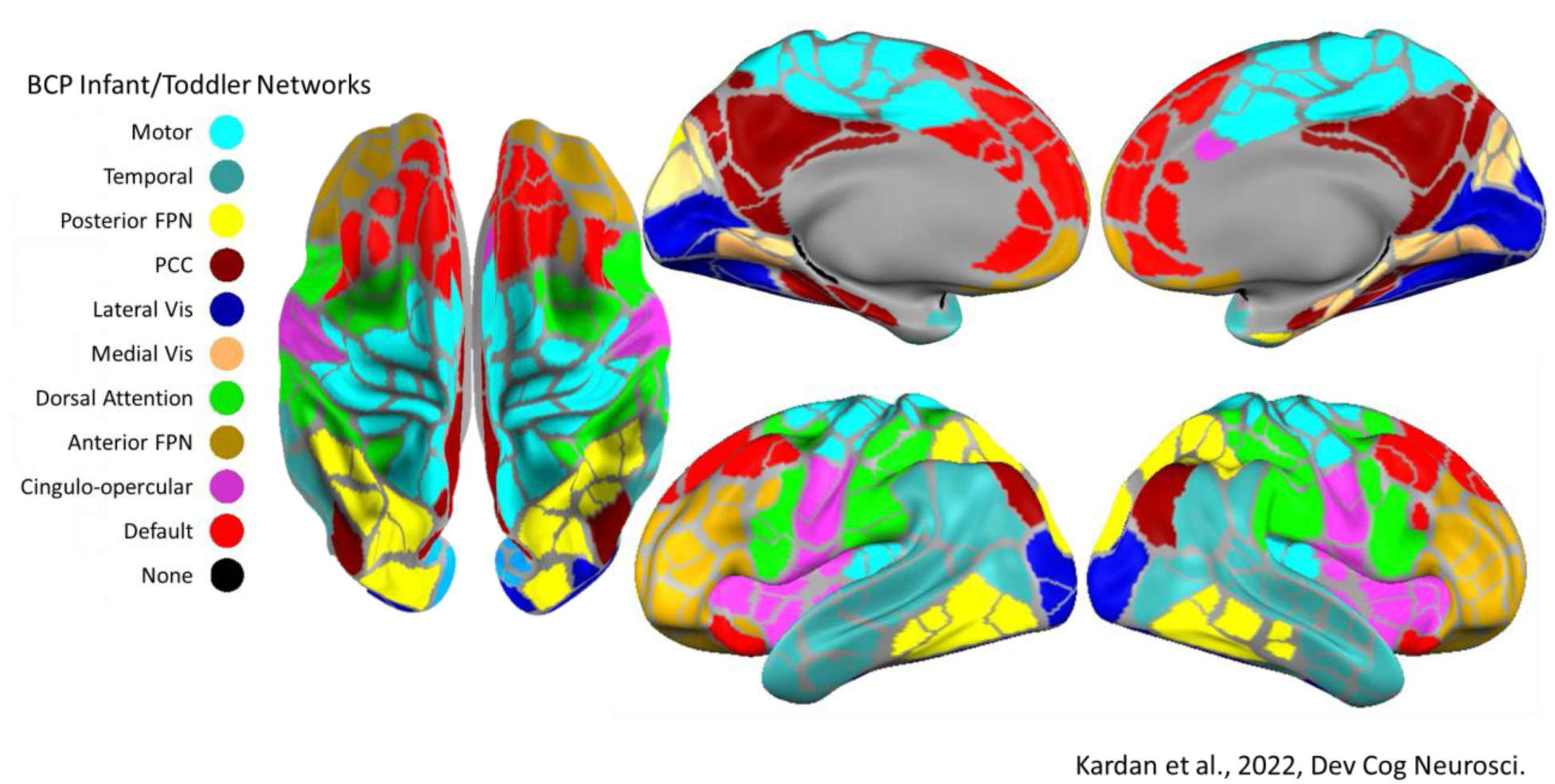
Baby network assignments. Note: FPN = Fronto-parietal network; PCC = Posterior cingulate cortex; Vis = Visual cortex; CO = Cingulo-opercular; DMN = Default Mode Network

**Figure S3.3.**
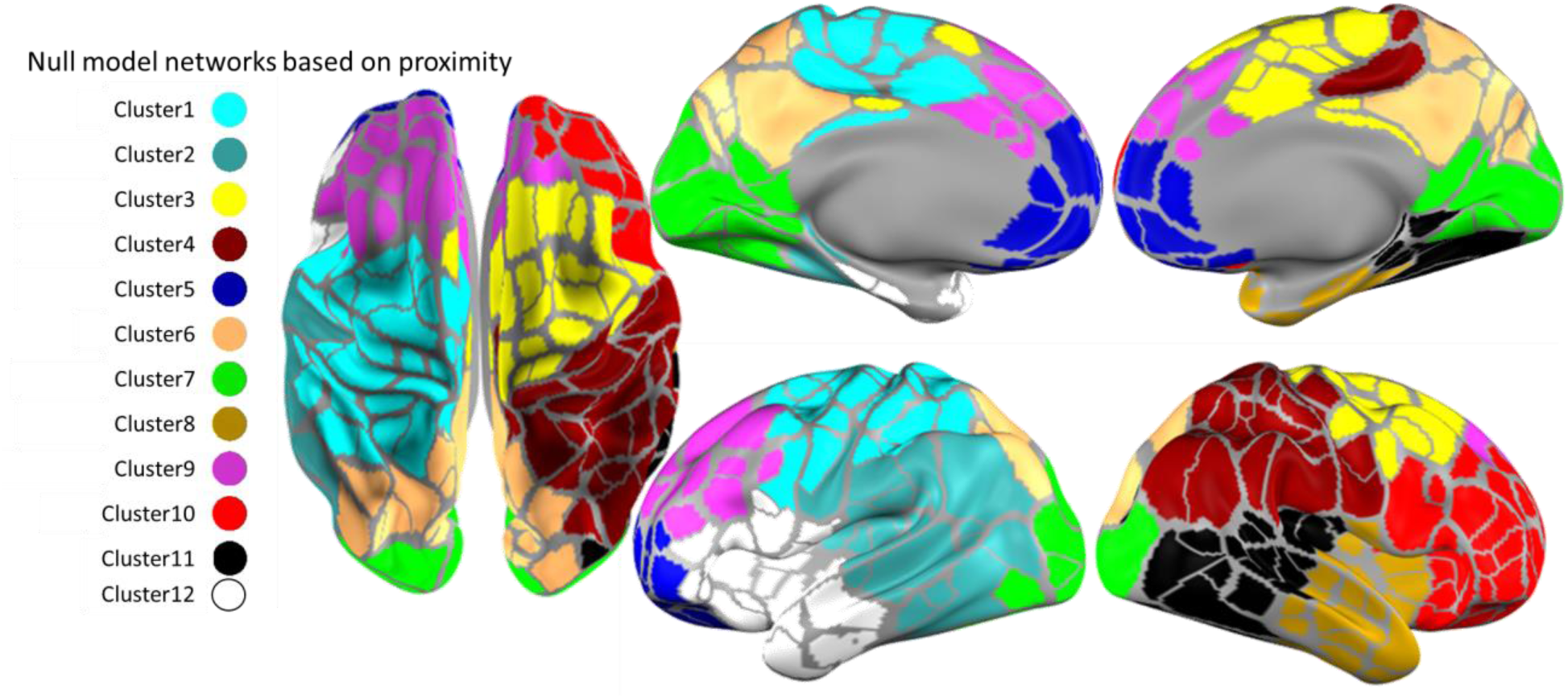
Proximity-based null networks based on k-means clustering on the parcel coordinates.

## Footnotes

1 Modularity can also be quantified for pre-determined communities instead of connectome-specific communities. If we take the difference in modularity to baby and adult networks to make anchored rsFC score, the resulting values would only have trivial differences with the current way of measuring the anchored rsFC score (i.e., based on Conn_within_ - Conn_between_), without any prediction gains for task performance.

## References

Avery, E. W., Yoo, K., Rosenberg, M. D., Greene, A. S., Gao, S., Na, D. L., … & Chun, M. M. (2020). Distributed patterns of functional connectivity predict working memory performance in novel healthy and memory-impaired individuals. Journal of cognitive neuroscience, 32(2), 241–255.

Ball, G., Kelly, C. E., Beare, R., & Seal, M. L. (2021). Individual variation underlying brain age estimates in typical development. Neuroimage, 235, 118036.

Bethlehem, R. A., Seidlitz, J., White, S. R., Vogel, J. W., Anderson, K. M., Adamson, C., … & Schaare, H. L. (2022). Brain charts for the human lifespan. Nature, 604(7906), 525–533.

Biondo, F., Jewell, A., Pritchard, M., Aarsland, D., Steves, C. J., Mueller, C., & Cole, J. H. (2022). Brain-age is associated with progression to dementia in memory clinic patients. NeuroImage: Clinical, 36, 103175.

Callaghan, B. L., & Tottenham, N. (2016). The stress acceleration hypothesis: Effects of early-life adversity on emotion circuits and behavior. Current opinion in behavioral sciences, 7, 76–81.

Casey, B. J., Cannonier, T., Conley, M. I., Cohen, A. O., Barch, D. M., Heitzeg, M. M., … & Dale, A. M. (2018). The adolescent brain cognitive development (ABCD) study: imaging acquisition across 21 sites. Developmental cognitive neuroscience, 32, 43–54.

Chan, M. Y., Na, J., Agres, P. F., Savalia, N. K., Park, D. C., & Wig, G. S. (2018). Socioeconomic status moderates age-related differences in the brain’s functional network organization and anatomy across the adult lifespan. Proceedings of the National Academy of Sciences, 115(22), E5144–E5153.

Chan, M. Y., Park, D. C., Savalia, N. K., Petersen, S. E., & Wig, G. S. (2014). Decreased segregation of brain systems across the healthy adult lifespan. Proceedings of the National Academy of Sciences, 111(46), E4997–E5006.

Chen, J., Tam, A., Kebets, V., Orban, C., Ooi, L. Q. R., Asplund, C. L., … & Yeo, B. T. (2022). Shared and unique brain network features predict cognitive, personality, and mental health scores in the ABCD study. Nature communications, 13(1), 1–17.

Cicchetti, D., & Toth, S. L. (2009). The past achievements and future promises of developmental psychopathology: The coming of age of a discipline. Journal of Child Psychology and Psychiatry, *50*(1-2), 16-25.

Cohen, J. R., & D’Esposito, M. (2016). The segregation and integration of distinct brain networks and their relationship to cognition. Journal of Neuroscience, 36(48), 12083–12094.

Corriveau, A., Yoo, K., Kwon, Y. H., Chun, M. M., & Rosenberg, M. D. (2023). Functional connectome stability and optimality are markers of cognitive performance. Cerebral Cortex, 33(8), 5025–5041.

Dehestani, N., Whittle, S., Vijayakumar, N., & Silk, T. J. (2023). Developmental brain changes during puberty and associations with mental health problems. Developmental Cognitive Neuroscience, 60, 101227.

Dong, H. M., Margulies, D. S., Zuo, X. N., & Holmes, A. J. (2021). Shifting gradients of macroscale cortical organization mark the transition from childhood to adolescence. Proceedings of the National Academy of Sciences, 118(28), e2024448118.

Eickhoff, S. B., Yeo, B. T. T., & Genon, S. (2018). Imaging-based parcellations of the human brain. Nature Reviews Neuroscience, 19(11), Article 11.

Eggebrecht, A. T., Elison, J. T., Feczko, E., Todorov, A., Wolff, J. J., Kandala, S., … & Pruett Jr, J. R. (2017). Joint attention and brain functional connectivity in infants and toddlers. Cerebral Cortex, 27(3), 1709–1720.

Erus, G., Battapady, H., Satterthwaite, T. D., Hakonarson, H., Gur, R. E., Davatzikos, C., & Gur, R. C. (2015). Imaging patterns of brain development and their relationship to cognition. Cerebral cortex, 25(6), 1676–1684.

Eyre, M., Fitzgibbon, S. P., Ciarrusta, J., Cordero-Grande, L., Price, A. N., Poppe, T., … & Edwards, A. D. (2021). The Developing Human Connectome Project: typical and disrupted perinatal functional connectivity. Brain, 144(7), 2199–2213.

Foulkes, L., & Blakemore, S. J. (2018). Studying individual differences in human adolescent brain development. Nature neuroscience, 21(3), 315–323.

Franke, K., Ziegler, G., Klöppel, S., Gaser, C., & Alzheimer’s Disease Neuroimaging Initiative. (2010). Estimating the age of healthy subjects from T1-weighted MRI scans using kernel methods: exploring the influence of various parameters. Neuroimage, 50(3), 883–892.

Fuhrmann, D., Knoll, L. J., & Blakemore, S. J. (2015). Adolescence as a sensitive period of brain development. Trends in cognitive sciences, 19(10), 558–566.

Funder, D. C., & Ozer, D. J. (2019). Evaluating effect size in psychological research: Sense and nonsense. Advances in methods and practices in psychological science, 2(2), 156–168.

Gao, S., Greene, A. S., Constable, R. T., & Scheinost, D. (2019). Combining multiple connectomes improves predictive modeling of phenotypic measures. NeuroImage, 201, 116038.

Gee, D. G. (2021). Early adversity and development: parsing heterogeneity and identifying pathways of risk and resilience. American Journal of Psychiatry, 178(11), 998–1013.

Gordon, E. M., Laumann, T. O., Adeyemo, B., Huckins, J. F., Kelley, W. M., & Petersen, S. E. (2016). Generation and evaluation of a cortical area parcellation from resting-state correlations. Cerebral cortex, 26(1), 288–303.

Hagler Jr, D. J., Hatton, S., Cornejo, M. D., Makowski, C., Fair, D. A., Dick, A. S., … & Dale, A. M. (2019). Image processing and analysis methods for the Adolescent Brain Cognitive Development Study. Neuroimage, 202, 116091.

Holm, M. C., Leonardsen, E. H., Beck, D., Dahl, A., Kjelkenes, R., de Lange, A. M. G., & Westlye, L. T. (2023). Linking brain maturation and puberty during early adolescence using longitudinal brain age prediction in the ABCD cohort. Developmental Cognitive Neuroscience, 60, 101220.

Howell, B. R., Styner, M. A., Gao, W., Yap, P. T., Wang, L., Baluyot, K., … & Elison, J. T. (2019). The UNC/UMN Baby Connectome Project (BCP): An overview of the study design and protocol development. NeuroImage, 185, 891–905.

Huys, Q. J., Maia, T. V., & Frank, M. J. (2016). Computational psychiatry as a bridge from neuroscience to clinical applications. Nature neuroscience, 19(3), 404–413.

Hyde, L. W., Bezek, J. L., & Michael, C. (2024). The future of neuroscience in developmental psychopathology. Development and Psychopathology, 1–16.

Johnson, S. B., Riis, J. L., & Noble, K. G. (2016). State of the art review: poverty and the developing brain. Pediatrics, 137(4).

Kardan, O., Stier, A. J., Cardenas-Iniguez, C., Schertz, K. E., Pruin, J. C., Deng, Y., … & Rosenberg, M. D. (2022). Differences in the functional brain architecture of sustained attention and working memory in youth and adults. PLoS Biology, 20(12), e3001938.

Kardan, O., Kaplan, S., Wheelock, M. D., Feczko, E., Day, T. K., Miranda-Domínguez, Ó., … & Rosenberg, M. D. (2022). Resting-state functional connectivity identifies individuals and predicts age in 8-to-26-month-olds. Developmental Cognitive Neuroscience, 56, 101123.

Lewis, J. D., Evans, A. C., Tohka, J., & Brain Development Cooperative Group. (2018). T1 white/gray contrast as a predictor of chronological age, and an index of cognitive performance. Neuroimage, 173, 341–350.

Maglanoc, L. A., Kaufmann, T., van der Meer, D., Marquand, A. F., Wolfers, T., Jonassen, R., … & Westlye, L. T. (2020). Brain connectome mapping of complex human traits and their polygenic architecture using machine learning. Biological psychiatry, 87(8), 717–726.

Marek, S., Tervo-Clemmens, B., Calabro, F. J., Montez, D. F., Kay, B. P., Hatoum, A. S., … & Dosenbach, N. U. (2022). Reproducible brain-wide association studies require thousands of individuals. Nature, 603(7902), 654–660.

Margulies, D. S., Ghosh, S. S., Goulas, A., Falkiewicz, M., Huntenburg, J. M., Langs, G., … & Smallwood, J. (2016). Situating the default-mode network along a principal gradient of macroscale cortical organization. Proceedings of the National Academy of Sciences, 113(44), 12574–12579.

Michael, C., Taxali, A., Angstadt, M., Kardan, O., Weigard, A., Molloy, M. F., … & Sripada, C. (2023). Socioeconomic resources in youth are linked to divergent patterns of network integration and segregation across the brain’s transmodal axis. bioRxiv.

Newman, M. E. (2006). Finding community structure in networks using the eigenvectors of matrices. *Physical Review E—Statistical*, Nonlinear, and Soft Matter Physics, 74(3), 036104.

Nielsen, A. N., Barch, D. M., Petersen, S. E., Schlaggar, B. L., & Greene, D. J. (2020). Machine learning with neuroimaging: evaluating its applications in psychiatry. Biological Psychiatry: Cognitive Neuroscience and Neuroimaging, 5(8), 791–798.

Owens, M. M., Potter, A., Hyatt, C. S., Albaugh, M., Thompson, W. K., Jernigan, T., … & Garavan, H. (2021). Recalibrating expectations about effect size: A multi-method survey of effect sizes in the ABCD study. PloS one, 16(9), e0257535.

Paus, T. (2005). Mapping brain maturation and cognitive development during adolescence. Trends in cognitive sciences, 9(2), 60–68.

Petrican, R., Taylor, M. J., & Grady, C. L. (2017). Trajectories of brain system maturation from childhood to older adulthood: Implications for lifespan cognitive functioning. Neuroimage, 163, 125–149.

Rosenberg, M. D., Finn, E. S., Scheinost, D., Papademetris, X., Shen, X., Constable, R. T., & Chun, M. M. (2016). A neuromarker of sustained attention from whole-brain functional connectivity. Nature neuroscience, 19(1), 165–171.

Rosvall, M., & Bergstrom, C. T. (2008). Maps of random walks on complex networks reveal community structure. Proceedings of the national academy of sciences, 105(4), 1118–1123.

Rubinov, M., & Sporns, O. (2010). Complex network measures of brain connectivity: uses and interpretations. Neuroimage, 52(3), 1059–1069.

Rudolph, M. D., Miranda-Domínguez, O., Cohen, A. O., Breiner, K., Steinberg, L., Bonnie, R. J.,… & Fair, D. A. (2017). At risk of being risky: The relationship between “brain age” under emotional states and risk preference. Developmental cognitive neuroscience, 24, 93–106.

Sripada, C., Rutherford, S., Angstadt, M., Thompson, W. K., Luciana, M., Weigard, A., … & Heitzeg, M. (2020). Prediction of neurocognition in youth from resting state fMRI. Molecular psychiatry, 25(12), 3413–3421.

Sripada, C., Angstadt, M., Taxali, A., Clark, D. A., Greathouse, T., Rutherford, S., … & Heitzeg, M. (2021). Brain-wide functional connectivity patterns support general cognitive ability and mediate effects of socioeconomic status in youth. Translational psychiatry, 11(1), 571.

Sylvester, C. M., Kaplan, S., Myers, M. J., Gordon, E. M., Schwarzlose, R. F., Alexopoulos, D., … & Smyser, C. D. (2023). Network-specific selectivity of functional connections in the neonatal brain. Cerebral Cortex, 33(5), 2200–2214.

Tooley, U. A., Bassett, D. S., & Mackey, A. P. (2021). Environmental influences on the pace of brain development. Nature Reviews Neuroscience, 22(6), 372–384.

Tooley, U. A., Bassett, D. S., & Mackey, A. P. (2022). Functional brain network community structure in childhood: Unfinished territories and fuzzy boundaries. NeuroImage, 247, 118843.

Wang, F., Zhang, H., Wu, Z., Hu, D., Zhou, Z., Girault, J. B., … & Li, G. (2023). Fine-grained functional parcellation maps of the infant cerebral cortex. elife, 12, e75401.

Whitmore, L. B., Weston, S. J., & Mills, K. L. (2023). BrainAGE as a measure of maturation during early adolescence. Imaging Neuroscience, 1, 1–21.

Xia, Y., Xia, M., Liu, J., Liao, X., Lei, T., Liang, X., … & He, Y. (2022). Development of functional connectome gradients during childhood and adolescence. Science bulletin, 67(10), 1049–1061.

Yarkoni, T., & Westfall, J. (2017). Choosing prediction over explanation in psychology: Lessons from machine learning. Perspectives on Psychological Science, 12(6), 1100–1122.

Yeung, A. W. K., More, S., Wu, J., & Eickhoff, S. B. (2022). Reporting details of neuroimaging studies on individual traits prediction: a literature survey. Neuroimage, 256, 119275.

Yoo, K., Rosenberg, M. D., Noble, S., Scheinost, D., Constable, R. T., & Chun, M. M. (2019). Multivariate approaches improve the reliability and validity of functional connectivity and prediction of individual behaviors. NeuroImage, 197, 212–223.

Zhang, Shaoshi, Bart Larsen, Valerie J. Sydnor, Tianchu Zeng, Lijun An, Xiaoxuan Yan, Ru Kong et al. “In vivo whole-cortex marker of excitation-inhibition ratio indexes cortical maturation and cognitive ability in youth.” Proceedings of the National Academy of Sciences 121, no. 23 (2024): e2318641121.

Zhao, X., Lynch Jr, J. G., & Chen, Q. (2010). Reconsidering Baron and Kenny: Myths and truths about mediation analysis. Journal of consumer research, 37(2), 197–206.

Zuo, X. N., He, Y., Betzel, R. F., Colcombe, S., Sporns, O., & Milham, M. P. (2017). Human connectomics across the life span. Trends in cognitive sciences, 21(1), 32–45.

